# Intermittent subdiffusion of short nuclear actin rods due to interactions with chromatin

**DOI:** 10.1101/2023.11.03.565456

**Authors:** Konstantin Speckner, Florian Rehfeldt, Matthias Weiss

## Abstract

The interior of cellular nuclei, the nucleoplasm, is a crowded fluid that is pervaded by protein-decorated DNA polymers, the chromatin. Due to the complex architecture of chromatin and a multitude of associated non-equilbrium processes, e.g. DNA repair, the nucleoplasm can be expected to feature non-trivial material properties and hence anomalous transport phenomena. Here, we have used single-particle tracking on nuclear actin rods, which are important players in DNA repair, to probe such transport phenomena. Our analysis reveals that short actin rods in the nucleus show an intermittent, anti-persistent subdiffusion with clear signatures of fractional Brownian motion. Moreover, the diffusive motion is heterogeneous with clear signatures of an intermittent switching of trajectories between at least two different mobilities, most likely due to transient associations with chromatin. In line with this interpretation, hyperosmotic stress is seen to stall the motion of nuclear actin rods, whereas hypoosmotic conditions yield a reptation-like motion. Our data highlight the local heterogeneity of the nucleoplasm, e.g. distinct biochemical microenvironments and chromosome territories, that need to be taken into account for an understanding of nucleoplasmic transport and the mechanobiology of nuclei.

## I. INTRODUCTION

Single-particle tracking (SPT) experiments have boosted our understanding of transport processes in soft and living matter (see [1–3] for reviews). Analyses of SPT data have shed light, for example, on the motion of lipids and proteins in membranes [4–7], on the diffusion of inert tracers in biomimetic fluids [8–10], or transport of mesoscopic complexes and organelles in the cytoplasm of living cells [11–15]. A frequent observation when analyzing SPT trajectories from complex and/or living systems is a sublinear scaling of the mean square displacement (MSD), commonly refered to as ‘subdiffusion’. Many elaborate models and stochastic processes have been discussed to explain the experimentally observed subdiffusive motion (see [16, 17] for an overview), starting from obstructed motion in a static, fractal maze of obstacles, via transient trapping and immobilization events with scale-free sojourn times, up to memory-containing random walks. The most widespread model for the latter class of subdiffusive transport processes is fractional Brownian motion (FBM) [18], a self-similar process with stationary, anti-correlated, and Gaussian increments that may be used as a clean mathematical model for the motion in viscoelastic media. Notably, standard Brownian motion is included in FBM as a special case in which the anti-persistent memory kernel approaches the Markovian limit.

Unlike fairly simple transport phenomena that only invoke a single stochastic process with fixed transport properties, e.g. FBM with a fixed memory kernel, many cases have been shown to feature a considerable diffusion heterogeneity, especially when dealing with living specimen. In these cases, step sizes taken by the tracked particle within a given period are not described by a simple Gaussian statistics but rather the distribution of step lengths is broadened and might even approach a Laplace distribution. Such a diffusion heterogeneity may arise from spatiotemporal fluctuations in the material properties of the medium explored by the tracked particle, which can be modeled by diffusing-diffusivity models [19–22]. Alternatively, an intermittent switching between two (or more) mobility states within single trajectories can account for the diffusion heterogeneity. Such an intermittent switching between at least two mobility states has been observed, for example, for RNA complexes in bacteria and yeast [11] and for quantum dots in the cytoplasm of living cells [14, 23, 24]. Furthermore, ambient non-equilibrium processes have been reported to contribute, for example, to the dynamics of chromatin [25–27] and telomeres [28, 29], suggesting that diffusion heterogeneity and/or intermittency might include or even rely on active processes.

Inspired by all of these findings, we wondered about the motion characteristics of short actin rods in the nucleus of living cells: Actin is an active filament that can propell itself in a ballistic fashion (’treadmilling’), it associates with chromatin strands at least when being involved in DNA damage repair [30, 31], and it is a non-spherical tracer that will explore a densly crowded nucleoplasm that is interspersed with polymeric chromatin segments. In fact, after it has been firmly established that the nucleus indeed harbors actin filaments (mostly short rods of some 10-100 nm length), transport properties of nuclear actin have not yet been quantified thoroughly. Therefore, a number of relevant questions have remained open: Does the motion of nuclear actin comprise active contributions, e.g. due to motor-driven DNA repair processes [32, 33], simple actin treadmilling [34], or due to an indirect shaking of the nucleoplasm by cytoplasmic microtubules [29]? How viscoelastic is the nucleoplasm for actin, is there an intermittent transport heterogeneity, and how much is the motion obstructed by decondensed chromatin?

To address these questions, we have performed SPT experiments on nuclear actin in transiently transfected culture cells and employed our recently established SPT analysis toolbox [35], in which also a thorough introduction to the relevant measures is given. As a result, we find that nuclear actin shows an intermittent, heterogeneous subdiffusion with an FBM-like memory and little active contributions in cells at physiological conditions. Upon exposing cells to osmotic stress, the intermittent diffusion behavior subsides, suggesting (transient) interactions with chromatin to be responsible for the diffusion heterogeneity. In particular, hyperosmotic conditions were seen to completely stall the motion of nuclear actin, whereas hypoosmotic stress leads to a reptation-like motion of nuclear actin that is in agreement with earlier observations with chromatin ends (telomeres). From these data, we conclude that the motion pattern of nuclear actin is strongly determined by the polymer dynamics of the chromatin.

## II. MATERIALS AND METHODS

### A. Cell culture and sample preparation

Bone osteosarcoma cells (U2OS, DSMZ Cat. # ACC-785, RRID: CVCL 0042) were cultured in McCoy’s 5A (Modified) Medium (# 26600023) supplemented with 10% fetal bovine serum (qualified, Brazil # 10270106), 1% L-glutamine (# A2916801), 1% sodium pyruvate (# 11360070) and 1% penicillin/streptomycin (# 15140122) (all from Gibco, Germany) in T-25 flasks (BioLite # 130189 Thermo Scientific, Germany) at 37°C and 5% CO_2_. Cells were split at 80% confluency every three days using pre-warmed trypsin/EDTA (0.25%) (# 25200056, Gibco, Germany) up to a maximum of 20 passages per cell batch. All culture cells were regularly checked by PCR to be free of mycoplasma contaminations.

For fluorescence microscopy, transiently transfected cells were plated on four-well *μ*-slides (ibidi, Germany). Nuclear actin was visualized by transient transfection with Utr230-EGFP-3XNLS [30] (Addgene plasmid # 58466, RRID: Addgene 58466) 24 h prior to microscopy using electroporation with the X-Unit of a 4D-Nucleofector device (Lonza Group, Switzerland) in SE Cell Line buffer using the program ‘CM-104’. Following the manufacturer’s protocol, the cell pellet was resuspended in 100 *μ*l transfection solution (82 *μ*l nucleofector solution + 18 *μ*l Supplement 1), transferred to a single nucleocuvette vessel containing 2 *μ*l plasmid DNA. Transfected cells were taken up in 500 *μ*l culture medium and seeded at a density of 75,000 cells per well. For coimaging of nuclei and actin, cells were transiently transfected 24 h prior to microscopy with H2B-mCherry [36] (Addgene plasmid # 20972, RRID: Addgene 20972) and Utr230-EGFP-3XNLS using Lipofectamine 3000 according to the manufactures protocol. Briefly, 400 ng plasmid DNA was incubated with 1 *μ*l P3000 and 1.5 *μ*l Lipofectamine 3000 reagent for 15 min in 50 *μ*l serum-free Opti-MEM (# 31985062, Gibco, Germany) and 35 *μ*l of the transfection solution was added dropwise to each well containing 40,000 cells.

For standard imaging, culture medium was substituted by MEM without phenol red, supplemented with 10% HEPES and 1% penicillin/streptomycin (referred to as ‘imaging medium’). For hypoosmotic conditions, the medium was replaced with 5% culture medium in 95% ultrapure water (Invitrogen, Waltham, Massachusetts, USA). During hyperosmotic treatment, cells were exposed to a solution of 1 M sucrose (Roth, Germany) in imaging medium. The cell medium was changed to the respective conditions 20 min prior to imaging in all cases. To depolymerize microtubules, cells were treated with 10 *μ*M nocodazole and transiently chilled on ice, as described and validated before [29]. In brief, a 2 mM stock solution of nocodazole (# M1404 Sigma Aldrich, Germany) dissolved in dimethylsulfoxide (# A3672 AppliChem, Germany) was diluted to working concentration in imaging medium. Nocodazole-treated cells were chilled on ice for 10 min before incubating at 37°C with 5% CO_2_ for 15 min. Imaging was performed subsequently in the presence of nocodazole.

### B. Microscopy, image analysis and tracking

Prior to imaging, cells were washed twice in Dulbecco’s phosphate-buffered saline (# 14190144 Gibco, Germany) and were subsequently supplemented with the corresponding imaging medium. Imaging was performed at 37°C with a customized spinning-disk confocal microscope, consisting of a Leica DMI 4000 microscope body (Leica Microsystems, Germany), a CSU-X1 (Yokogawa, Japan) spinning disk unit, and a custom-made incubation chamber. Images were acquired using a Hamamatsu Orca Flash 4V2.0 sCMOS camera (Hamamatsu, Japan) paired with an HC PL APO 63x/1.4 (Leica Microsystems) oil immersion objective, resulting in a pixel size of 56.2 nm. Illumination of the specimen at 490 nm and 561 nm was achieved by solid-state lasers (Calypso and Jive, Cobolt, Sweden), the corresponding fluorescence was detected by bandpass filters (Semrock, USA) in the range 500-550 nm and 575-625 nm. The whole setup was controlled by custom written LabView software (National Instruments, USA). For single-particle tracking of nuclear actin, time-resolved image stacks were taken at an interval of 100 ms (up to 1250 images in total) using a 2 × 2 pixel binning. For two-color imaging of nuclear actin and chromatin, z-stacks with a separation of 200 nm and 35 confocal layers were acquired using an exposure time of 200 ms without binning. During measurements, the ventilation of the preheated air-conditioning chamber was suspended.

Fluorescence images were imported by FIJI [37] and processed for subsequent tracking of nuclear actin. First, images were cropped to only include the cell area and the residual background was subtracted. Next, contrast was enhanced by enlarging the fluorescence images fourfold and reducing image noise below two pixels with a band-pass filter. The time-series images were scaled down by a factor of two and subjected to drift correction via the ImageJ/FIJI plugin Stackreg [38]. Particle positions were detected and linked to trajectories by the ImageJ/FIJI plugin TrackMate (version 6.0.3) [39]. As an input parameter for TrackMate, the diameter of nuclear actin filaments was estimated from their intensity profiles, yielding a tracking diameter of 350 ± 50 nm. Particle tracking was performed using the Laplacian-of-Gaussian (LoG) detection algorithm (threshold set to 2000 ±500 and subpixel localization). Spots touching the nuclear envelope were removed and positions were linked with the simple linear assignment problem (LAP) tracker. Here, a maximum linking distance of 300 nm was used and gaps of three frames, not exceding more than 5% of all detections, were allowed per trajectory. The minimum trajectory length was set to 50 positions and non-assigned detections were cleaned from the time series. Typically, about 50 localizations were found in a single image with about 250 trajectories obtained from individual nuclei.

Particle trajectories were exported as XML files and converted to ASCII files for further processing in Matlab (Matlab 2018b, The MathWorks Inc., USA). In our analyses, trajectories of nuclear actin were truncated to *N* = 100 time steps, i.e. shorter trajectories were discarded. In total, our ensembles consisted of 1500-5000 trajectories of at least ten cell nuclei (depending on the particular condition, ensemble sizes stated in the main text).

## III. RESULTS AND DISCUSSION

Being a central player in virtually all aspects of mechanobiology, one can expect filamentous actin to also be involved in many dynamical processes of the nucleus. In line with this notion, the nucleus softens, for example, after inducing DNA damage [40] and short nuclear actin filaments have been shown to be involved in repair of such damages [30, 31]. Based on this interaction with chromatin and since the nucleus has a complex mechanobiology [41], dynamic transport processes of actin filaments in the nucleoplasm can be expected to have intricate features. Some of the open questions in this context have been phrased already in the Introduction.

To gain insights into these aspects, we have performed and analyzed SPT experiments on nuclear actin in transiently transfected U2OS cells (see Materials and Methods for details). In particular, we have stained actin by UTR-GFP, for which long and persistent fibers at the cell cortex and short rods in the nucleoplasm were expected [30] and observed (see Fig. 1a for representative examples). Due to the diffraction limit, short actin rods in the nucleus appeared point-like and were hence ideal for tracking experiments. Retaining only trajectories with at least *N* = 100 positions (longer ones were trimmed to this length), we used our recently introduced toolbox [35] to explore and analyze the mode of motion of nuclear actin in detail.

**FIG. 1.**
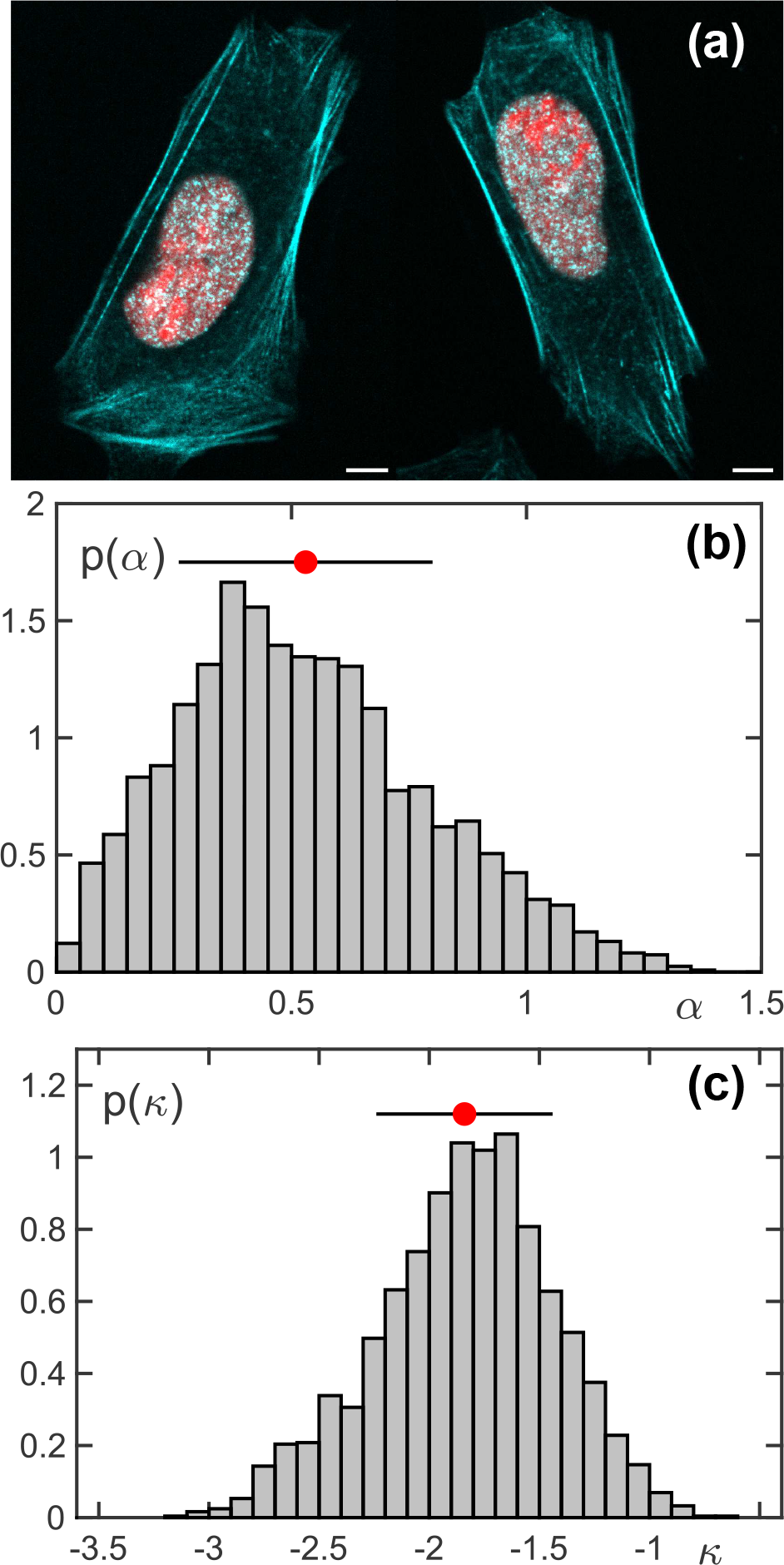
(a) Representative images of U2OS cells in which actin was visualized by UTR-GFP (cyan) and histones marked by H2B-mCherry (red). Long actin stress fibers are visible especially at the cell cortex whereas short nuclear actin rods appear point-like due to the diffraction limit; scale bars 5 *μ*m. (b) The PDF of scaling exponents, *p*(*α*), obtained from fitting TA-MSDs of all trajectories in untreated cells, features a mean ⟨*α* ⟩= 0.53 (red circle) at a considerable standard deviation (black horizontal bar), representing the variation between individual trajectories. Hence, nuclear actin rods move, on average, clearly subdiffusive. (c) The associated PDF of generalized diffusion coefficients, here reported as dimensionless quantity *κ* = log_10_(*K* · 1 *s*^*α*^ */μ*m^2^), features an almost lognormal shape, i.e. the area explored on the 1 s-time scale varies widely between individual trajectories (mean and standard deviation shown as red circle and black bar).

As a first and basic step, we have calculated for each trajectory the time-averaged mean square displacement (TA-MSD). The TA-MSD is supposedly the simplest meaningful quantity that can be extracted for individual trajectories, providing first insights into the spreading behavior of individual actin rods. For our ensemble of *M* = 2452 trajectories in untreated cells (positions indexed in integer multiples of the frame time Δ*t* = 100 s), the TA-MSD for each trajectory reads

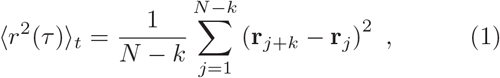

where *τ* = *k*Δ*t* denotes the lag time. Since MSDs (and also the velocity autcorrelation function, see below) indicated a negligible influence of localization errors, we have fitted all TA-MSDs in the range 3Δ*t* ≤ *τ* ≤ 15Δ*t* with a simple power law of the form

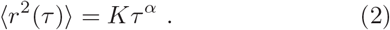

Here *K* denotes the generalized diffusion coefficient, while the scaling exponent *α* captures deviations from the linear scaling of normal Brownian motion.

The resulting probability distribution function (PDF) *p*(*α*) featured a mean ⟨*α* ⟩ = 0.53, which clearly indicates that nuclear actin shows, on average, a marked subdiffusion (Fig. 1b). The considerable width of *p*(*α*), i.e. the standard deviation *σ* ≈ 0.27 of the PDF, is not just due to statistical errors that arise from limited averaging in short trajectories [42] but rather reflects the variation in the motion of individual trajectories that all explore and report on their own local environments. Interestingly, only a fraction of about 1% of all trajectories featured a scaling exponent *α* ≥ 1.2, i.e. the SPT data strongly suggest that nuclear actin shows a purely diffusive rather than a ballistic transport.

For inspecting the associated PDF of generalized diffusion coefficients, we have chosen to define a dimensionless quantity *κ* = log_10_(*K* · 1*s*^*α*^/*μ*m^2^), since values for *K* not only varied widely between individual trajectories (due to varying local environments) but also since values for *K* are not directly comparable for different values of *α*. Values of *κ* therefore report the logarithm of the typical area (in *μ*m^2^) that is explored within one second, indicating the rapidity of the random motion on the one-second time scale. As a result, we observed *p*(*κ*) to feature a roughly lognormal shape (Fig. 1c) that reflects the broad variation of local mobilities of nuclear actin rods in their respective environment (see also discussion in [10] on the emergence of the lognormal shape).

Having revealed that nuclear actin rods do not show ballistic signatures but rather are subject to a marked subdiffusion, we next probed whether there is a memory associated with this erratic motion. To this end, we calculated the ensemble average of all trajectory-wise velocity autocorrelation functions (VACFs), defined as

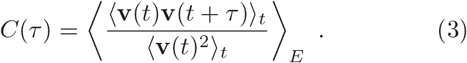

Here, *τ* = *k*Δ*t* denotes the lag time and v_*j*_ = [r*_j+n_* − r_*j*_]*/*δ*t* is the instantaneous velocity, determined by steps r*_j+n_* − r_*j*_ taken within an integer multiple of the frame time, *δt* = *n*Δ*t*. For probing the impact of localization errors and for comparison with analytical expressions, it is convenient to inspect the VACF as a function of the normalized time *ξ* = *τ/δt*. As can be seen in Fig. 2a, the VACFs for different choices of *δt* fall on top of each other, yielding a single master curve with a pronounced negative minimum, irrespective of the value of *δt*. The negative minimum indicates a significant anti-persistent memory in the trajectories and the congruence at *ξ* = 1 for all choices of *δt* highlights that the anti-correlation is not an artificial and transient effect of localization errors [43]. Moreover, the data is in very good agreement with an analytical expression for FBM,

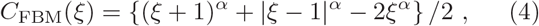

suggesting that the observed subdiffusion of nuclear actin can be described by an anti-persistent FBM process. Notably, the value *α* = 0.6 that needs to be used in Eq. (4) to match the experimental data is slightly higher than the mean of *p*(*α*) (cf. Fig. 1b), which might not be overly surprising when bearing in mind the considerable width of the PDF.

**FIG. 2.**
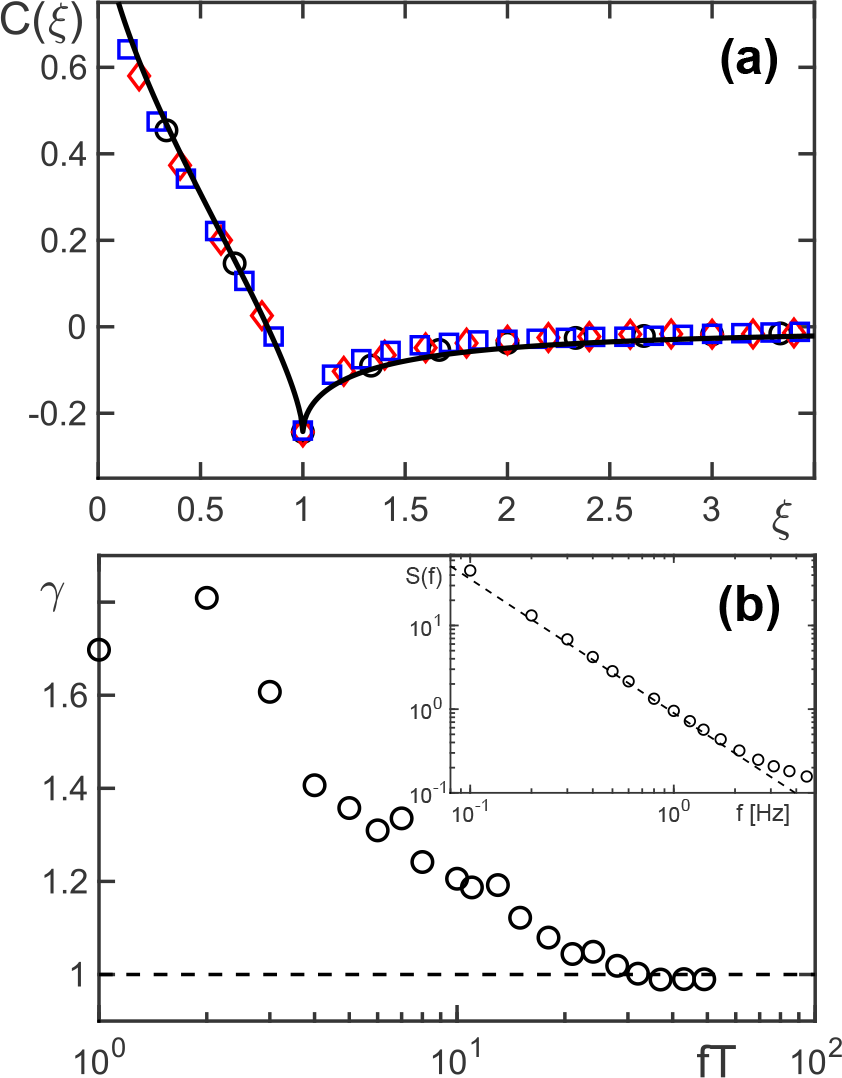
(a) The ensemble-averaged VACFs, *C*(*ξ*), obtained for periods *δt/*Δ*t* = 3, 5, 7 (black circles, red diamonds, blue squares), all coincide when rescaling lag times as *ξ* = *τ/δt*. This observation indicates that localization errors are negligible and that the random walk of nuclear actin features a considerable anti-persistent memory. The very good agreement with the FBM prediction [Eq. (4)] (black line, *α* = 0.6) suggests that nuclear actin exhibits an anti-persistent subdiffusion that is typically observed in viscoelastic media. (b) Analyzing the associated PSDs provides further support for this interpretation: The ensemble-averaged PSD (inset, black circles) shows a power-law decay *S*(*f*)1*/f* ^1+*α*^ (black dashed line, *α* = 0.6), as anticipated for FBM. The coefficient of variation, *γ*, for trajectory-wise PSDs with respect to their ensemble average converges to unity for large frequencies (main plot, dimensionless frequency *f T* = 1, *…, N*). This confirms that nuclear actin movement can be described as a subdiffusive FBM random walk.

To further probe the hypothesis that nuclear actin is fueled by an FBM process, we have calculated the power-spectral density (PSD)

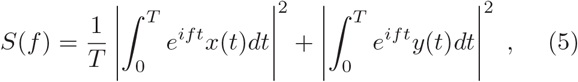

for each trajectory as well as the resulting ensemble-average. In line with previous predictions for subdiffusive FBM, individual and ensemble-averaged PSDs followed a power-law decay *S*(*f*) ∼ 1*/f* ^1+*α*^ (Fig. 2b, inset). Moreover, individual PSDs showed fluctuations around the ensemble average that are captured by the PSDs’ coefficient of variation,

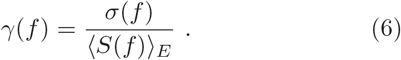

For subdiffusive FBM processes, an asymtotic convergence *γ* → 1 has been predicted and verified even in the presence of localization errors [44, 45]. The data for nuclear actin nicely follow this very prediction (see Fig. 2b), providing additional support to the hypothesis that the motion can be captured by an anti-persistent FBM random walk. Notably, these results are in agreement with a previous assessment of the motion of nuclear actin [30] but go significantly deeper in the analysis as more measures are employed to extract and quantify the type of random walk.

Summarizing the results obtained for untreated cells until here, we have found that nuclear actin rods show an anti-correlated subdiffusion that is consistent with an anti-persistent FBM random walk process. No signatures for an active, ballistic motion were detected, i.e. if treadmilling occurs for nuclear actin, it is invisible on the probed length and time scales. The average scaling exponent *α* ≈ 0.55±0.05 (found from MSDs, VACFs, and PSDs), is surprisingly similar to those found for individual telomeres [29, 46] but also for the motion of inert tracers [47] and biomolecular condensates [48] in the nucleoplasm. This suggests that the anti-persistent dynamics of nuclear actin emerges due to the same, polymer-physics related processes [49]: Assuming that chromatin strands can be modeled as simple Rouse polymers, the motion of a monomer in such a polymer (e.g. a telomere) will show subdiffusion of the FBM type with *α* = 1*/*2 for short and intermediate time scales (i.e. up to the Rouse time scale). Therefore, if short nuclear actin rods (transiently) associate with chromatin, they can be expected to also move like an effective monomer, consistent with the data shown in Figs. 1 and 2. The existence of a (transient) association with chromatin is corroborated by the involvement of nuclear actin in DNA repair processes [30, 31]. A second interpretation, albeit similar in spirit, uses somewhat larger length scales, i.e. an ensemble of Rouse polymers, for which the emerging medium is known to be viscoelastic with a scaling of the complex shear modulus as |*G*(*ω*)| ∼ *ω*^1*/*2^, hence enforcing an FBM random walk of tracer particles, again with *α* = 1*/*2. In this scenario, the viscoelastic medium surrounding actin filaments enforces an FBM random walk with very similar features as for the aforementioned monomer motion, but supposedly with a different (and elevated) diffusion coefficient. Either way, the subdiffusion observed for nuclear actin may be traced back qualitatively and quantitatively to chromatin’s polymer physics and the subsequent analysis will provide hints which of the two interpretations is more likely.

To follow up on this, we have probed whether trajectories are governed by a single Gaussian process, as expected for a simple FBM random walk, or if rather a marked diffusion heterogeneity is observed for nuclear actin. To this end, we have extracted for each trajectory the one-dimensional steps in *x*- and *y*-direction, taken within a period *δt*, and normalized these sets of increments by their respective root-mean-square value to make trajectories comparable (see [35] for a more detailed discussion). With this approach, the mean step length is fixed to unity and the PDF of all normalized increments *p*(*χ*) will comply with a normal distribution for any *δt* unless a diffusion heterogeneity within individual trajectories is present. Since we did neither observe differences between the spatial directions nor between positive and negative steps, we combined all normalized steps into a single PDF of step moduli, *p*(|*χ*|), sometimes refered to as van-Hove function [16]. As a result, we observed significant deviations from a normal distribution for small-scale steps, i.e. when choosing *δt* = Δ*t* (Fig. 3a). The marked deviations from a simple Gaussian highlight a considerable diffusion heterogeneity, i.e. nuclear actin appears to switch between at least two different modes of motion within single trajectories. In fact, the experimental data were well described by a superposition of two Gaussians. Only when analyzing steps on larger scales, i.e. when choosing *δt* = 20Δ*t*, the experimental data were in reasonable agreement with a normal distribution (Fig. 3a). Similar observations have been made before for tracers in the cytoplasm [14] and they were attributed to transient associations with the endomembrane system, leading to an intermittent change of the diffusion coefficient.

**FIG. 3.**
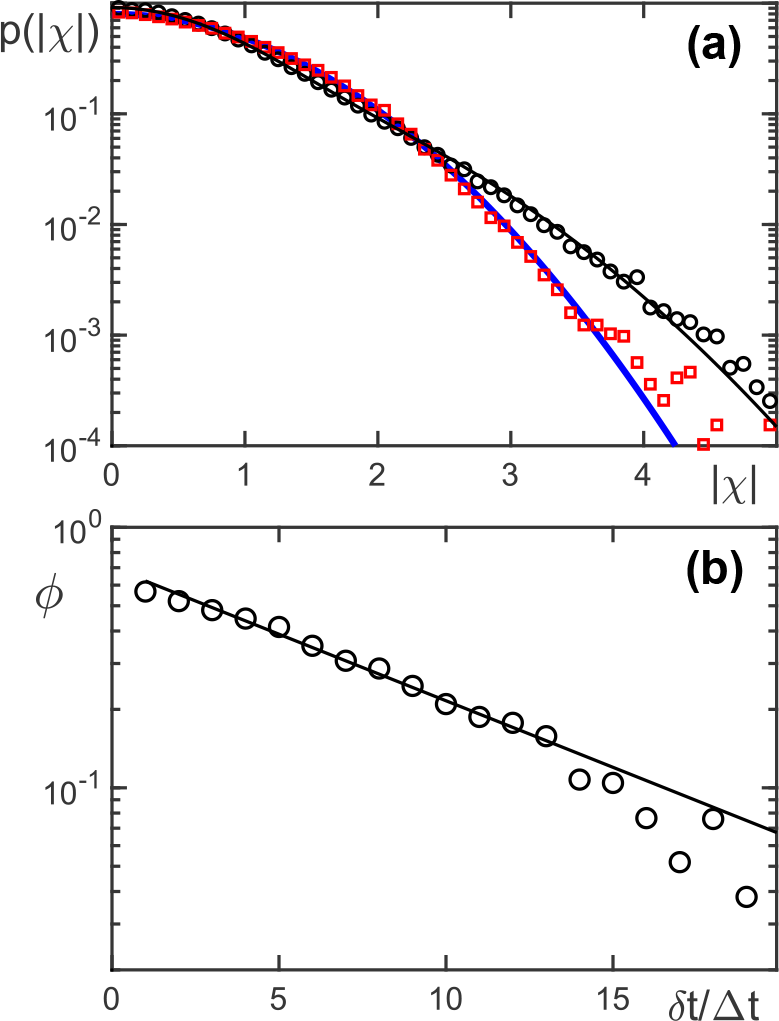
(a) The PDF of normalized steps taken within a period *δt*, here shown as *p*(|*χ*|), strongly deviates from a normal distribution (blue line) for *δt* = Δ*t* (black circles) but is well captured by a superposition of two Gaussians (black line). In contrast, for *δt* = 20Δ*t* (red squares) a resonable agreement with a single Gaussian is observed. (b) The fraction *ϕ* (black circles), contributed to *p*(|*χ*|) by a Gaussian process with lower mobility, decreases for increasing periods *δt* in an approximately exponential fashion (black line).

To probe such an intermittent character of the dynamics of nuclear actin, we have evaluated the deviations of *p*(|*χ*|) from a single Gaussian as a function of the period needed for the step, *δt*. Given that a superposition of only two Gaussians provided already a very good fit for *p*(|*χ*|) (cf. Fig. 3a), we have modeled the PDFs of normalized step moduli for all *δt* with a superposition of only two Gaussians,

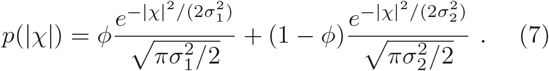

Asking for the relative contributions of two Gaussian processes with different average step length, we reasoned that the ratio 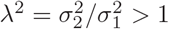 has to remain constant for all *δt*, whereas *ϕ* (the contribution of the random walk with smaller steps, i.e. a lower mobility) should be an open parameter. Moreover, since Eq. (7) is used to fit a PDF of experimentally obtained and already normalized increments, the two variances are actually coupled and can be expressed as *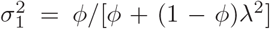* and *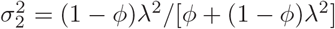*.

As a result of fitting *p*(|*χ*|) for several choices of *δt* in the range |*χ*| *<* 4 (for which statistics was sufficient in all cases), we have found a very good description of the experimental data for *λ*^2^ = 3.4 and *ϕ* ≈ 0.7 exp(−*δt/*(8.5Δ*t*)) (Fig. 3b). In other words, the motion of nuclear actin on long time scales is well described by a single Gaussian random walk process, whereas for small and intermediate time scales the contribution of a second Gaussian process needs to be considered. The diffusive mobility encoded in the two Gaussians differs approximately threefold. In particular, on the 1 s time scale, i.e. for *δt* ≈ 10Δ*t*, the slower process has almost died out and only a single Gaussian process with higher mobility contributes to the motion of actin rods. An interpretation for this intermittent behavior could be the transient association of actin with chromatin, leading to a slow FBM-like motion that mimics the dynamics of a monomer in a Rouse polymer, followed by a more rapid motion in the viscoelastic nucleoplasm (without an explicit linking to individual chromatin segments) after detaching from chromatin on the 1 s time scale. It is hence conceivable that actin behaves intermittently like a monomer in a Rouse chain before dissociating and exploring the viscoelastic surrounding in a free manner. This interpretation will be tested below by removing the free voids between chromatin strands via osmotic pressure.

Given that telomeres, i.e. monomers of chromatin, have been shown to be driven by external active noise, mediated by microtubules that shake the nucleus [29], we next aimed at exploring how much the intermittent subdiffusion of nuclear actin rods is actively driven. To this end, we treated cells with nocodazole to disrupt microtubules (see Materials and Methods) and repeated the SPT experiments (yielding an ensemble of *M* = 1672 trajectories). As a result of the analysis, we observed that in the absence of functional microtubules the gross behavior of the random walk barely changed, i.e. the PDF *p*(*α*) did not change significantly (as tested by a two-sample Kolmogorov-Smirnov test). This finding is also in agreement with the visual impression obtained from box plots for *α* (Fig. 4a). In contrast, the box plots for *κ* indicated a significant reduction, that translates to a roughly 1.4-fold reduction of the area that is explored within one second. Albeit this is less than the roughly twofold mobility change observed for telomeres under the same treatment [29], it clearly demonstrates that also nuclear actin is partially driven by an active, microtubule-mediated noise. Moreover, bearing in mind that nuclear actin rods move intermittently, with supposedly only the less mobile state being linked to chromatin, a 1.4-fold reduction of the generalized diffusion coefficient appears reasonable.

**FIG. 4.**
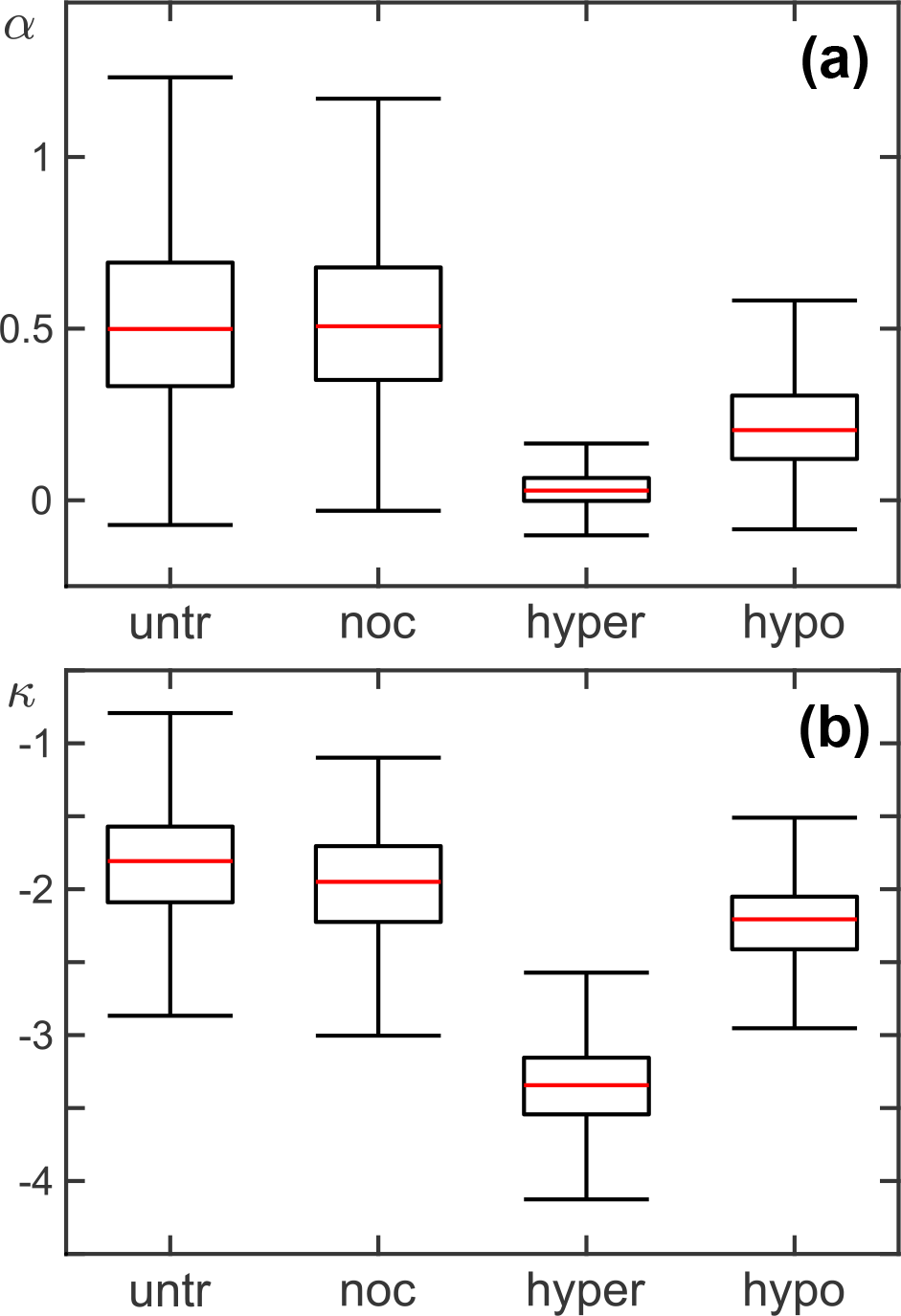
Boxplots for (a) the MSD scaling exponents, *α*, and (b) the generalized diffusion coefficients (again reported as dimensionless quantity *κ*), obtained for nuclear actin in untreated cells (’untr’), after nocodazole treatment (’noc’), upon applying hyperosmotic (’hyper’), or hypoosmotic stress (’hypo’). Pairwise comparison of the data by a two-sample Kolmogorov-Smirnov test (significance level *p* ≤ 0.001) revealed signifcant differences in all cases except for comparing *α*-values in untreated and nocodazole-treated cells. While *α* in the presence of nocodazole was not significantly altered with respect to cells at physiological conditions, highly significant changes were observed when applying osmotic stress: Hyperosmotic stress yielded an almost complete arrest of actin movement, whereas hypoosmotic stress resulted in a value *α* ≈ 1*/*4. Also data for *κ* at all four conditions were deemed significantly different, albeit the reduction of the area that is explored within one second, i.e. *K* · 1*s*^*α*^/*μ*m^2^, was reduced only 1.4-fold and 1.8-fold upon nocodazole treatment and application of hypoosmotic stress, respectively.

Given that the interpretation of SPT data of nuclear actin rely on arguments from polymer physics, we next aimed at changing the polymeric state of chromatin. To this end we applied osmotic stress to the cells since water loss or uptake can be expected to collapse or expand the polymeric chromatin. Here, ensembles of *M* = 5411 and *M* = 2289 trajectories were analyzed for hyper- and hypoosmotic conditions, respectively.

Applying hyperosmotic stress by adding sucrose to the cell medium had a drastic effect on the motion of nuclear actin with *α* approaching zero and *κ* decreasing to very small values (Fig. 4). Therefore, the motion of nuclear actin was basically stalled in this case, in line with earlier reports on an irreversible collapse of chromatin to a dense, molten-globule-like configuration [50] and an almost complete immobilization of telomeres [46] at the same condition. In contrast, diluting the cell medium to apply hypoosmotic stress (see Materials and Methods), nuclear actin rods remained mobile but changed their key features significantly (Fig. 4): The scaling exponent *α* as well as the mobility parameter *κ* were reduced approximately twofold and no signatures for an intermittency were visible any more in *p*(|*χ*|); nuclear actin rods showed only one mode of motion. Interestingly, very similar values for *α* and *κ* had been found earlier for the motion of telomeres in hypoosmotically stressed cells [46], suggesting that the short nuclear actin rods are subject to the very same dynamics, namely that of a monomer within the swollen chromatin polymer. Since hypoosmotic stress induces a massive water uptake, leading to swollen cells and nuclei, chromatin can be expected to decondense and fill the nucleus in a disordered and entangled phenotype. Assuming chromatin to still be described by the Rouse model, the motion of a monomer below the Rouse time is predicted via the tube model to exhibit a MSD scaling exponent *α* = 1*/*4 [49, 51] which is in very nice agreement with the experimental observation (cf. Fig. 4). Given that no intermittent switching of mobilities is observed in this condition, actin rods would stay associated with chromatin (as an effective monomer) for fairly long periods. This interpretation is also in line with our conclusions for untreated cells (see above): A free motion at high mobility in voids between chromatin strands becomes suppressed by the swelling of chromatin, hence erradicating the contribution of a second Gaussian with larger step lengt in *p*(|*χ*|). The remaining chromatin-associated state, i.e. the slow motion like a monomer in the chromatin polymer, survives but gets updated to a reptation mode with lower *α*.

Taken together, our SPT data and their detailed analysis have revealed that short actin rods in the nucleus show an intermittent, anti-persistent subdiffusion with clear signatures of an FBM process with little contributions from an external active noise. Altering the state of chromatin by applying osmotic stress has revealed, that the motion of nuclear actin rods most likely reflects the dynamics of an effective monomer in a Rouse chain, switching stochastically forth and back to a free motion in the viscoelastic nucleoplasm. The broad variations between individual trajectories most likely reflects the heterogeneity experienced in the nucleus, e.g. arising from biochemically distinct microenvironments and persistent chromosome territories. The data shown here will hopefully facilitate subsequent analyses of other transport transport processes in the nucleoplasm as well as investigations on the complex mechanobiology of cell nuclei, especially in response to DNA damages.

## AUTHOR CONTRIBUTIONS

KS carried out all experiments, retrieved particle trajectories from image series, and performed initial analyses; FR and MW designed and supervised the study; MW wrote all Matlab codes and analyzed the data; all authors contributed to writing of the manuscript.

## ACKNOWLEDGEMENTS

Financial support by the VolkswagenStiftung (Az. 92738) and by the Elite Network of Bavaria (Study Program Biological Physics) are gratefully acknowledged.

## Notes

### Competing Interest Statement

The authors have declared no competing interest.

